# FBXW7 triggers degradation of WDR5 to prevent mitotic slippage

**DOI:** 10.1101/2022.08.24.505075

**Authors:** Simon Hänle-Kreidler, Kai T. Richter, Ingrid Hoffmann

## Abstract

During prolonged mitotic arrest induced by anti-microtubule drugs, cell fate decision is determined by two alternative pathways, one leading to cell death, the other inducing premature escape from mitosis by mitotic slippage. FBWX7, a member of the F-box family of proteins and substrate-targeting subunit of the SCF (SKP1-CUL1-F-Box) E3 ubiquitin ligase complex promotes mitotic cell death and prevents mitotic slippage. In this study, we report that WDR5, a component of the mixed lineage leukemia (MLL) complex of Histone 3 Lysine 4 (H3K4) methyltransferases is a substrate of FBXW7. WDR5 binds to FBXW7 in vivo and in vitro and its ubiquitin-mediated proteasomal degradation is mediated by FBXW7. Furthermore, we find that WDR5 depletion counteracts FBXW7 loss-of-function by reducing mitotic slippage and polyploidization. Our data elucidate a new mechanism in mitotic cell fate regulation which might contribute to prevent chemotherapy resistance in patients after anti-microtubule drug treatment.

## Introduction

FBXW7/hCdc4 is a substrate specificity component of the SCF (Skp1-Cullin-F-box E3 ubiquitin ligase) and a member of the F-box protein family which is responsible for recognizing and binding phosphorylated substrates to regulate their turnover (1). Most of the FBXW7 substrates are widely studied protooncogenes including Cyclin E, c-Myc, Notch and c-Jun (2). FBXW7 functions as tumor suppressor and mutations in the FBXW7 gene have been observed in a variety of human cancers (3) (2).

Several lines of evidence have implicated loss or mutation of FBXW7 in chemotherapy resistance against anti-microtubule drugs (4) (5). For example, taxanes are microtubule-stabilizing drugs that inhibit the dynamic instability of mitotic spindles and taxane-based chemotherapy agents are used for treatment of a wide range of cancers. Anti-microtubule drugs cause a prolonged mitotic delay and mitotic cell death. Perturbation of microtubule dynamics induces activation of the spindle assembly checkpoint (SAC) and mitotic arrest. However cells can escape from the SAC-induced mitotic arrest through a process called mitotic slippage (6). Upon mitotic arrest caused by anti-microtubule drugs, FBXW7 promotes cell death and prevents mitotic slippage (7) and is counteracted by the E3 ligase FBXO45/MYCBP2, a regulator of FBXW7 protein abundance in mitosis (8). Two competing cellular networks, cyclin B1 degradation and activation of proapoptotic caspases, were proposed to regulate cell fate following mitotic arrest (9). Mcl-1, a pro-survival member of the Bcl-2 family central to the intrinsic apoptosis pathway, is degraded during a prolonged mitotic arrest and may therefore act as a mitotic death timer (5). It is being debated whether FBXW7 has a direct role in regulating Mcl-1 protein levels (5) (10) (11) and identification of FBXW7 substrates in mitotic cell fate regulation is a major task.

WDR5 (WD-repeat containing protein 5) is a highly conserved core scaffolding subunit of the KMT2(MLL/SET) enzymes that deposit histone H3 lysine 4 (H3K4) methylation (12) (13) (14) (15). Apart from regulating gene expression by chromatin remodeling in interphase, WDR5 also functions in cell division (16) (17). In particular, WDR5 localizes to the midbody and regulates abscission (18). Moreover, WDR5 also regulates localization of the kinesin motor protein Kif2A during mitosis to facilitate chromosome congression and proper spindle assembly (19). In this study we identified WDR5 as a new substrate of SCF-FBXW7. FBXW7 ubiquitinates WDR5 and its depletion leads to increased WDR5 protein levels. Furthermore, we find that WDR5 promotes mitotic slippage. Our study also has translational implication to provide a rationale for the FBXW7/WDR5 axis as a possible target for chemotherapy resistance treatment in combination with anti-microtubule drug therapies.

## Results

### Proteomics-based approach to identify putative substrates of FBXW7

We have previously shown that the SCF-FBXW7 complex is targeted for degradation by the FBXO45-MYCBP2 E3 ubiquitin ligase specifically during mitotic arrest induced by anti-microtubule drugs. This in turn leads to induction of mitotic slippage and prevention of mitotic cell death (8). Now we aimed at identifying ubiquitination substrates of FBXW7 in mitotic slippage induction. We performed a screen with the aim to search for FBXW7 substrates that function in mitotic cell fate regulation (Fig. 1A). FLAG-tagged FBXW7α was immunopurified from HEK293T and analyzed by mass spectrometry. As a negative control, we used a FLAG-tagged FBXW7α-WD40 mutant that lacks the ability to bind substrates, but not Skp1 and Cul1 (20). Among the proteins that we identified by mass-spectrometry were c-Myc and members of the MLL-complex including KMT2D, WDR5 and ASH2L (Fig. 1A and Supplemental Fig. 1). To verify the findings of our screen, we transfected Flag-tagged FBXW7 in HEK-293T cells. Immunoprecipitation using anti-Flag affinity beads identified WDR5 and KMT2D (MLL4) as putative FBXW7 binding proteins. We find that the interaction between FBXW7 is likely specific for KMT2D as no interaction with two other MLL-family members SetD1A and MLL1 was detected (Figure 1B). The interaction between FBXW7 and KMT2D, as well as WDR5, was abolished when the WD40 domain of FBXW7 that includes the conserved amino acid residues R465, R479, and R505, required for high affinity substrate binding, was mutated (FBXW7^ARG^) (21). An interaction between FBXW7 and KMT2D was previously also identified by (22). As our major aim was to identify FBXW7 substrates involved in mitotic slippage, we first checked whether KMT2D could act as a FBXW7 substrate in mitotic slippage. We analyzed this by depleting KMT2D in U2OS cells and performed live-cell imaging in the presence of nocodazole. Our results shown in Fig. 1C reveal that KMT2D does not function in mitotic slippage.

**Figure 1.**
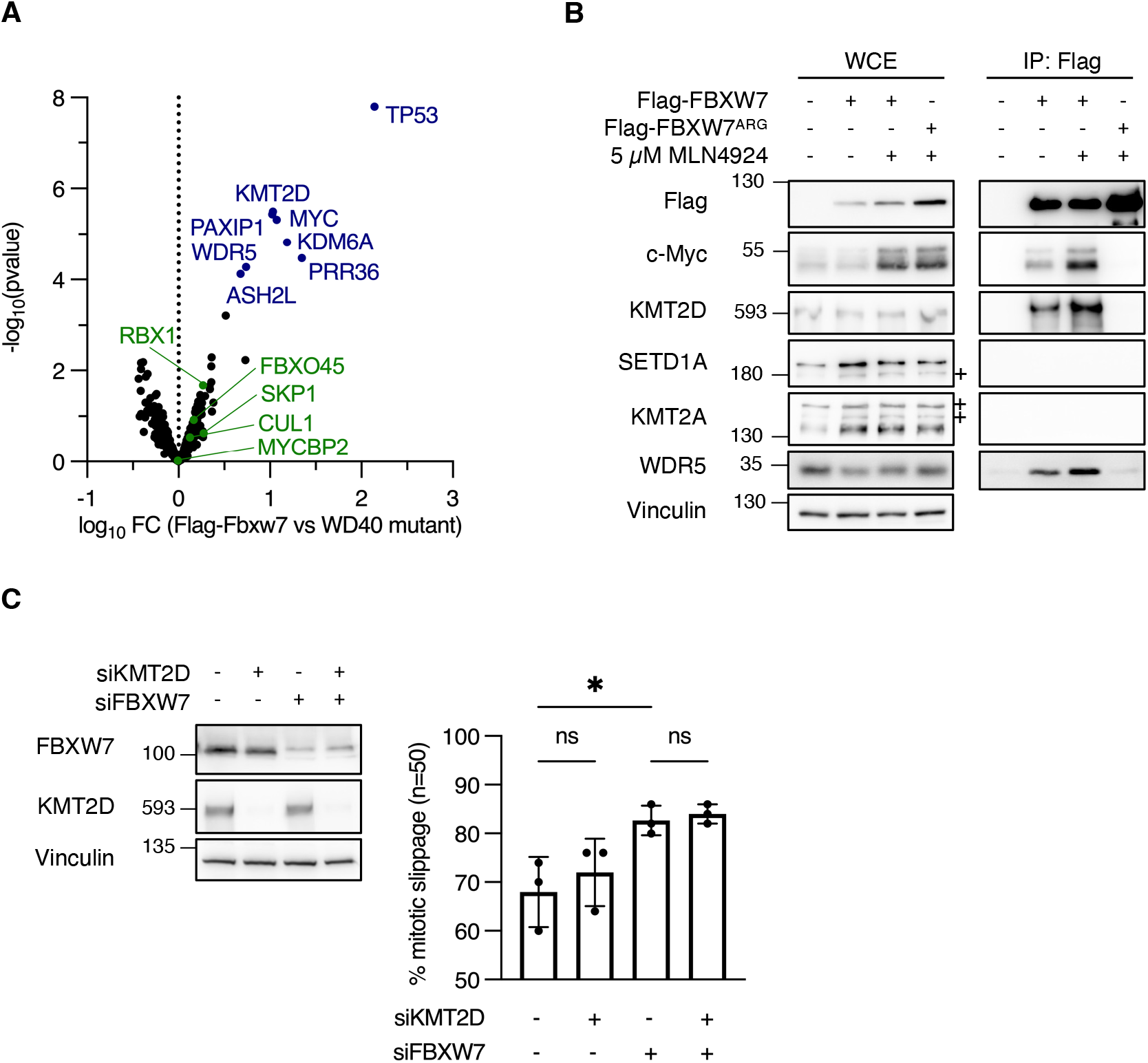
Proteomics based approach to identify putative FBXW7 substrates. A, volcano plot showing enrichment of FBXW7-WD40 interactors identified by IP-MS. Flag-FBXW7 or Flag-FBXW7 (T439I, S462A, T463A, R465A) were overexpressed in HEK293T for 48 h and interactors identified by Flag-immunoprecipitation, TMT10-plex labeling and mass spectrometry. Significance was determined via t-statistics using LIMMA R-package; proteins were considered significant if they showed a fold change of at least 50% and an adjusted p-value after Benjamini and Hochberg below 5%, n = 2. B, Flag-FBXW7 or Flag-FBXW7^ARG^ (R465H, R479Q, R505C mutations) were overexpressed in HEK293T cells for 48 h with or without inhibition of cullin-RING E3 ligases by 5 µM MLN-4924 for 5 h prior to harvest, as indicated. Flag-FBXW7 constructs were used for an immunoprecipitation against the FLAG-Tag. + marks non-specific bands. C, U2OS were transfected twice with 30 nM siRNA targeting GL2, KMT2D or FBXW7, as indicated. 830 nM nocodazole were added 48 h after the second siRNA transfection and mitotic cell fates of n = 50 cells determined by live-cell imaging. * p < 0.05, ns p > 0.05, one-way ANOVA with Tukey post-hoc test, n = 3.

### FBXW7 regulates WDR5 protein levels by ubiquitination and proteasomal degradation

It was previously shown that MLL regulates M-phase progression in association with the WRAD complex and a conserved subunit of WRAD, WDR5, localizes to the mitotic spindle and to the midbody in dividing human cells (19).

Interestingly, WDR5 was identified as a major interacting protein of FBXW7 in our screen and protein levels of WDR5 were decreased in response to FBXW7 overexpression (Fig. 1B). We therefore checked whether WDR5 protein levels would be affected by FBXW7 ablation. We found that in two FBXW7 knockout (KO) cell lines HCT116 and DLD1, as well as in HeLa cells transfected with siFBXW7, WDR5 protein levels were significantly increased (Fig. 2A), which might suggest that FBXW7 could also regulate WDR5 protein stability. qPCR experiments indicated that WDR mRNA levels remain unchanged and that WDR5 is upregulated at the post-translational level (Fig. 2B). WDR5 protein levels were also upregulated in FBXW7 KO cells, whether or not KMT2D was present, showing that WDR5 upregulation is independent of KMT2D (Fig. 2C). A phenylalanine residue within the WIN site of WDR5 (F133A) is required for the interaction with KMT2 methyltransferases (23). We asked whether FBXW7 could still bind to the WDR5-F133A mutant and therefore independently of KMT2D. Our results shown in Fig. 2D showed that this mutant can still co-precipitate FBXW7 but not KMT2D. To show that the interaction between FBXW7 and WDR5 is direct, we performed in vitro binding assays where we incubated insect cell-derived His-FBXW7/Skp1 with bacterially expressed GST-WDR5 and performed GST-pulldown assays. Our results presented in Fig. 2E show that FBXW7 directly binds to WDR5.

**Figure 2.**
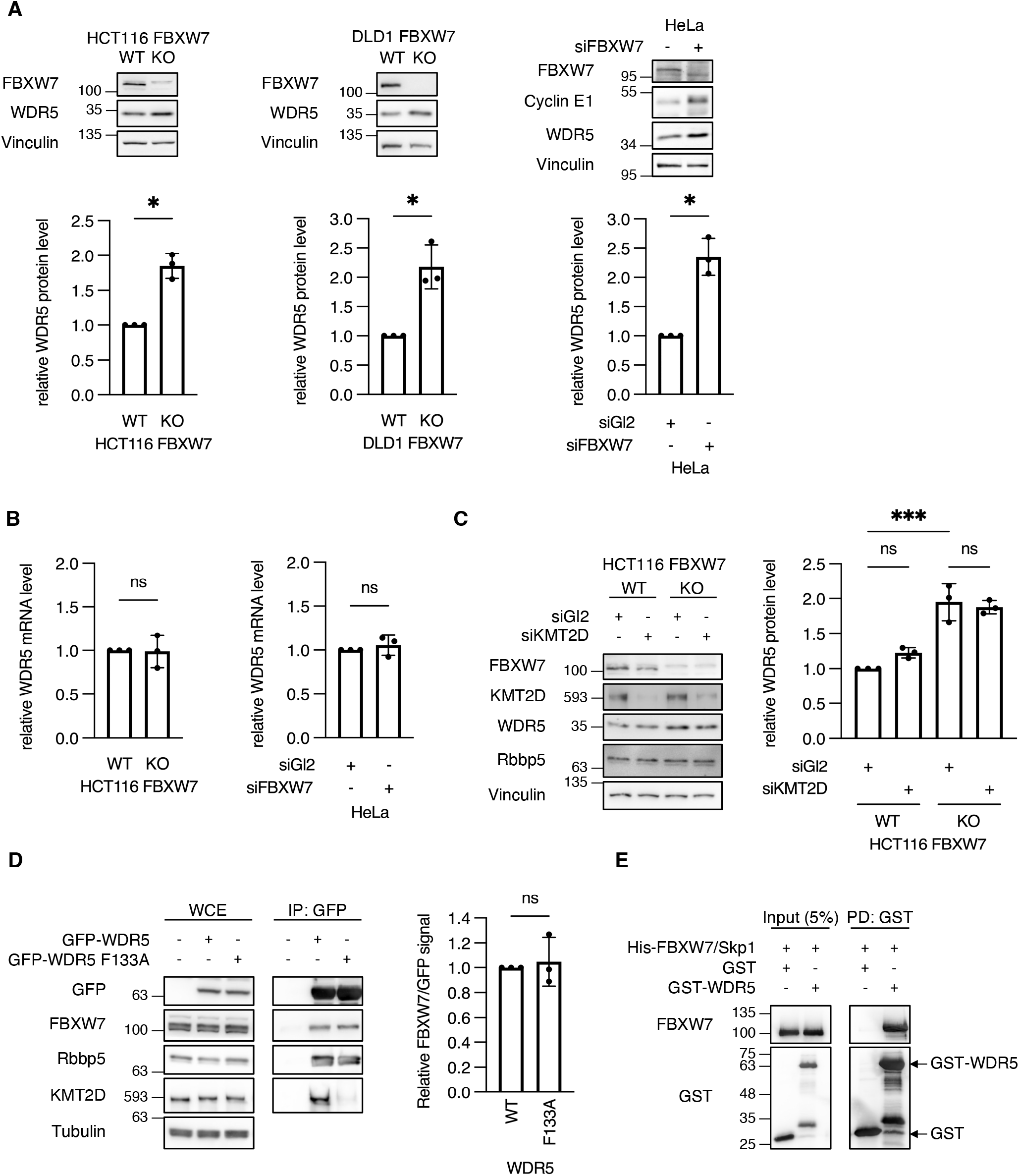
WDR5 is upregulated by FBXW7 depletion and interacts with FBXW7 independently of KMT2D. A, HeLa were transfected twice with siRNA against Gl2 or FBXW7 and harvested 72 h after the first transfection. * p-value < 0.05, unpaired two-way student’s t-test with Welch’s correction, n = 3. B, HeLa were transfected twice with siRNA against Gl2 or FBXW7 and harvested 72 h after the first transfection. ns p-value > 0.05, unpaired two-way student’s t-test with Welch’s correction, n = 3. C, HCT116 WT and FBXW7 knockout cell lines were transfected twice with siRNA against Gl2 or KMT2D and harvested 72 h after the first transfection. *** p-value < 0.001, ns p-value > 0.05, one-way ANOVA with Tukey post-hoc test, n = 3. D, GFP-WDR5 WT or a F133A mutant were overexpressed in HEK293T cells for 24 h and immunoprecipitated using GFP-trap beads. ns p-value > 0.05, unpaired two-way student’s t-test with Welch’s correction, n =3. E, purified recombinant GST-WDR5 and His-FBXW7 were combined and GST-pulldowns performed using reduced glutathione on CL-4B beads.

In order to assess the effect of FBXW7 on WDR5 protein stability, we used FBXW7 WT and KO DLD1 cell lines and added cycloheximide (CHX) to block translation. Cells were harvested at different time points after CHX addition. In FBXW7-depleted cells, overexpressed Flag-WDR5 (Fig. 3A) and endogenous WDR5 (Fig. 3B) were stabilized compared to FBXW7-wild-type cells (Fig. 3A), indicating that FBXW7 promotes the degradation of WDR5. To address whether WDR5 protein levels are regulated by proteasomal degradation, we treated cells with the proteasomal inhibitor MG132. In the presence of MG132, the decay of WDR5 was prevented (Fig. 3C). To analyze whether the regulation of WDR5 by FBXW7 is mediated by ubiquitination, we performed in vivo ubiquitination assays. We found that overexpression of FBXW7leads to an increase of WDR5 polyubiquitination (Fig. 3D lane 3 versus lane 2). Taken together, our findings suggest that WDR5 is a new substrate of FBXW7 which regulates its ubiquitin-dependent proteasomal degradation.

**Figure 3.**
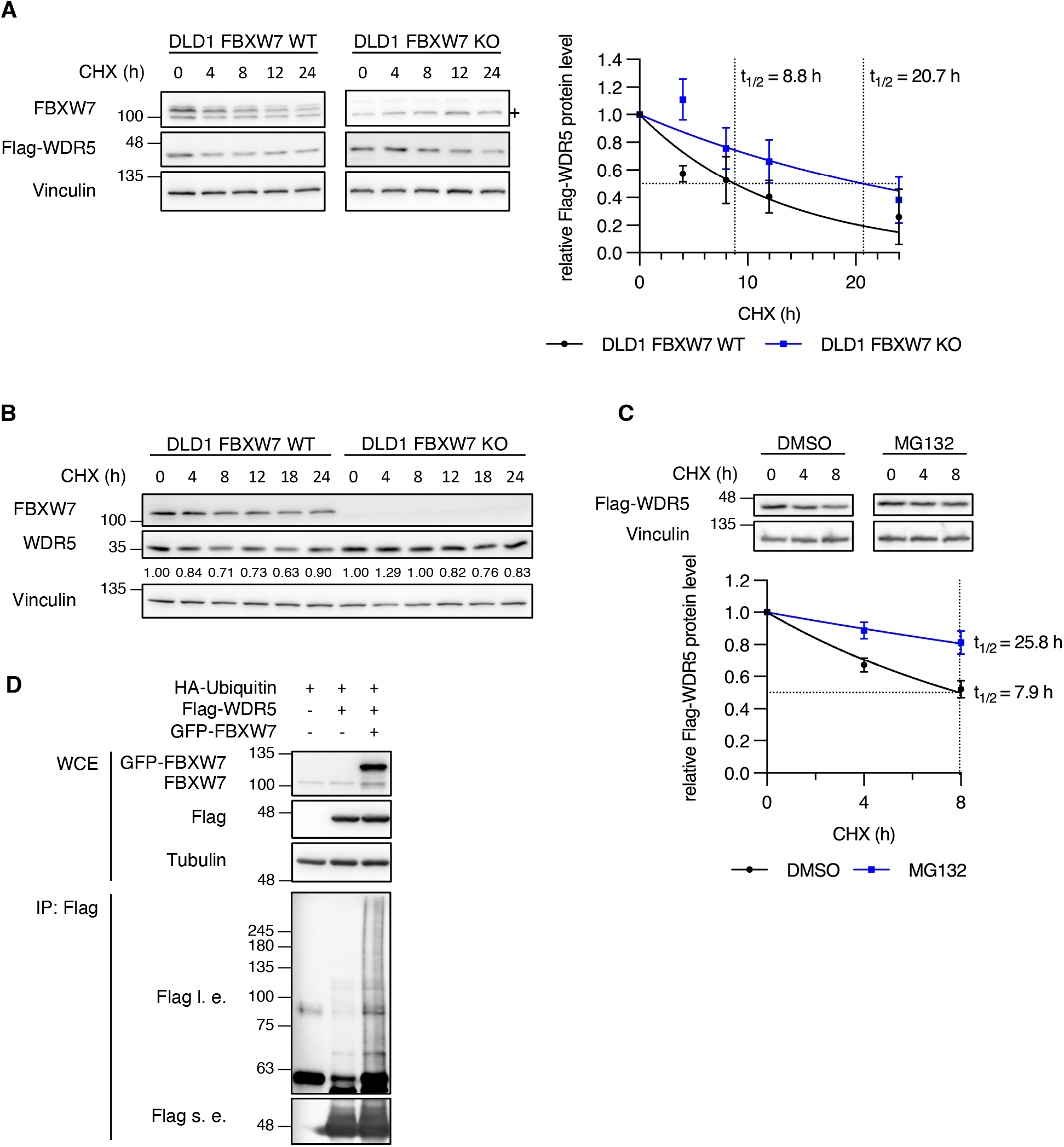
WDR5 protein stability is regulated by FBXW7-mediated ubiquitination. A, Flag-WDR5 was overexpressed in DLD1 WT and FBXW7 knockout cell lines for 24 h, followed by blockage of protein synthesis with 300 µg/ml cycloheximide for the indicated durations prior to harvest. Protein half-lives were determined by fitting to a one-phase decay model, n = 3. + marks a non-specific band. B, DLD1 WT and FBXW7 knockout cell lines were treated with 300 µg/ml cycloheximide for the indicated durations prior to harvest. C, Flag-WDR5 was overexpressed in DLD1 WT for 24 h, followed by inhibition of protein synthesis with 300 µg/ml cycloheximide in the presence of 10 µM MG132 or DMSO for the indicated durations prior to harvest. Normalized protein levels were fit to a one-phase decay model to determine protein half-lives, n = 3. D, the indicated proteins were overexpressed in HEK293T for 24 h and the 26S proteasome was blocked by 10 µM MG132 for 5 h prior to harvest. Flag-WDR5 was immunoprecipitated using α-Flag M2 beads in the presence of 20 mM N-ethylmaleimide. l. e. - long exposure, s. e. - short exposure.

The interaction between FBXW7 and WDR5 is partially regulated by GSK3ß-dependent phosphorylation Degradation of FBXW7 substrates depends on phosphorylation of a phosphodegron (2). To find out whether the interaction between FBXW7 and WDR5 is regulated by phosphorylation, we treated Flag-FBXW7 immunoprecipitates with λ-phosphatase and observed that the interaction with WDR5 was impaired (Fig. 4A). In a variety of substrates including Cyclin E, c-Myc and c-Jun, GSK3β phosphorylation plays a pivotal role (24) (25) (26). We therefore analyzed whether GSK3β-dependent phosphorylation of WDR5 is also required for the interaction between FBXW7 and WDR5. We made use of a small-molecule inhibitor, CHIR-99021, that blocks the enzymatic activity of GSK3β. Treatment of WDR5 overexpressing HEK293T cells with the GSK3β inhibitor reduced but did not completely block the binding between FBXW7 and WDR5 (Fig. 4B). Moreover, we found that siRNA-mediated ablation of GSK3β leads to upregulation of WDR5 protein levels (Fig. 4C). Co-expression of WDR5 and GSK3β followed by immunoprecipitation of WDR5 revealed an interaction of WDR5 with both ectopically expressed and endogenous GSK3β (Fig. 4D). We then analyzed whether treatment of HEK293T cells with CHIR-99021 would affect WDR5 ubiquitination. While WDR5 was ubiquitinated by FBXW7 in the control, this increase in ubiquitination was markedly reduced in the presence of the GSK3β inhibitor, as well as by inhibition of cullin-RING E3 ubiquitin ligases using the small molecule neddylation inhibitor, MLN4924 (Fig. 4E). Together, these results propose that GSK3β cooperates with FBXW7 to regulate WDR5 protein abundance.

**Figure 4.**
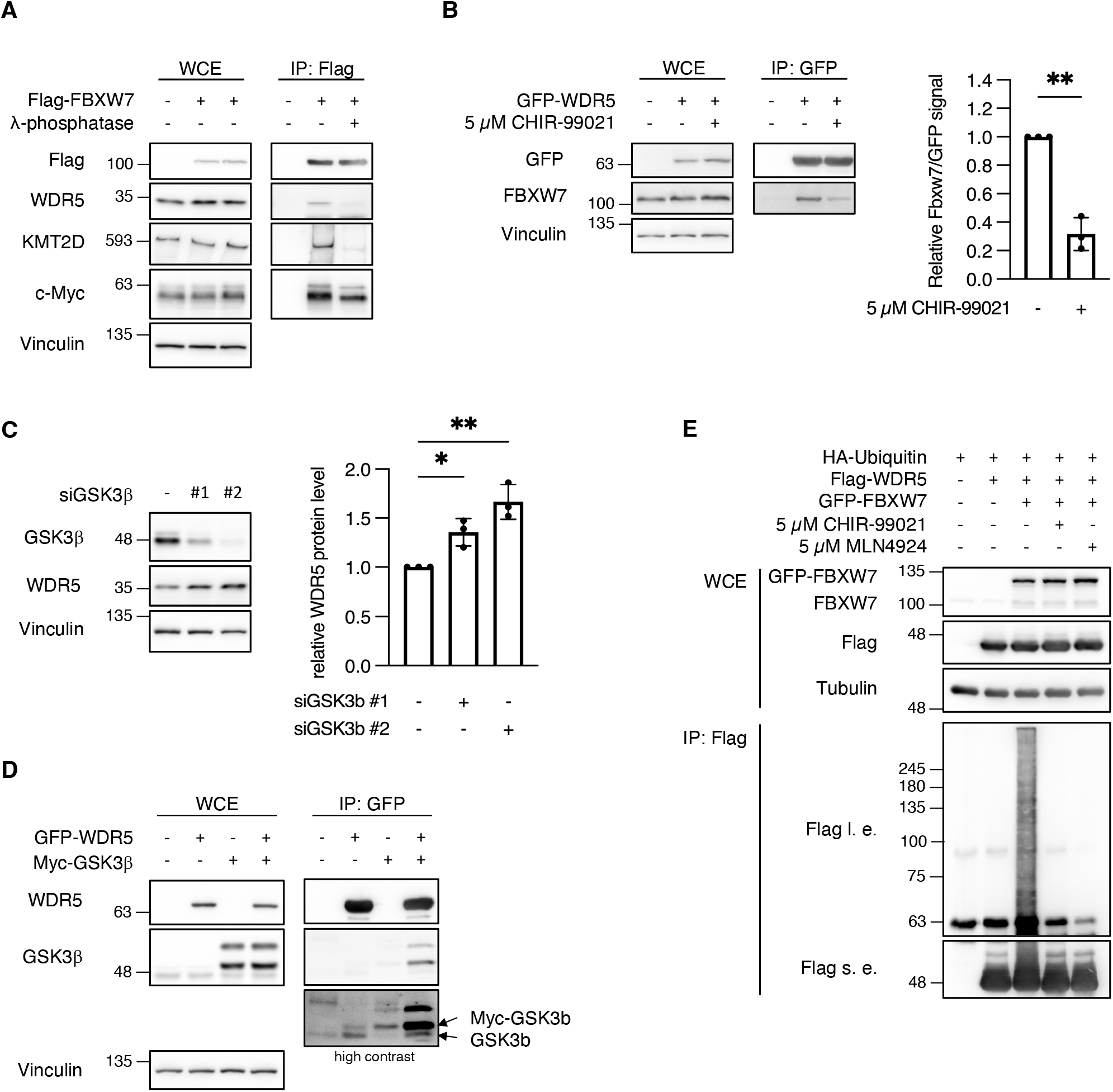
The regulation of WDR5 by FBXW7 depends in part on GSK3β. A, Flag-FBXW7 was overexpressed in HEK293T for 24 h and all cultures were treated with 5 µM MLN-4924 for 5 h prior to harvest. Flag-FBXW7 was immunoprecipitated and the indicated sample was dephosphorylated with λ-phosphatase at 30°C for 3 h. B, GFP-WDR5 was overexpressed in HEK293T for 24 h and cultures were treated with 5 µM CHIR-99021 or DMSO for 5 h prior to cell harvest, as indicated. GFP-WDR5 was immunoprecipitated using GFP-trap beads. ** p < 0.01, unpaired two-way student’s t-test with Welch’s correction, n = 3. C, U2OS cells were transfected twice with 30 nM against GL2 or GSK3β and harvested 72 h after the first transfection. * p < 0.05, ** p < 0.01, one-way ANOVA, n = 3. D, GFP-WDR5 and Myc-GSK3β were overexpressed in HEK293T cells for 24 h. GFP-WDR5 was immunoprecipitated using GFP-trap beads. E, the indicated proteins were overexpressed in HEK293T for 24 h. 5 µM CHIR-99021 or 5 µM MLN4924 were added for 9 h and 10 µM MG132 for 5 h prior to harvest. Flag-WDR5 was immunoprecipitated using α-Flag M2 beads in the presence of 20 mM N-ethylmaleimide. l. e. - long exposure, s. e. - short exposure.

WDR5 and Cyclin E1 promote mitotic slippage Previous reports indicate that FBXW7 prevents mitotic slippage by promoting mitotic cell death in cells that were arrested in mitosis upon treatment with anti-microtubule drugs (4) (5). In addition, (27) (19) showed that MLL and WDR5 regulate M-phase progression. Given the role of WDR5 in mitosis and the fact FBXW7 depletion leads to an upregulation of WDR5 protein levels (22) and Figure 2, we asked whether overexpression of WDR5 would influence mitotic slippage. We generated an U2OS cell line where WDR5 could be ectopically induced by doxycycline addition. We analyzed mitotic cell fate by live-cell imaging (Figure 5A, Videos 1 and 2) in the presence of spindle poisons at concentrations that prevent cell division and caused either mitotic cell death or mitotic slippage (28). Our results show that upregulation of WDR5 leads to an increase in mitotic slippage in cells that were arrested in mitosis with nocodazole (Figure 5B).

**Figure 5.**
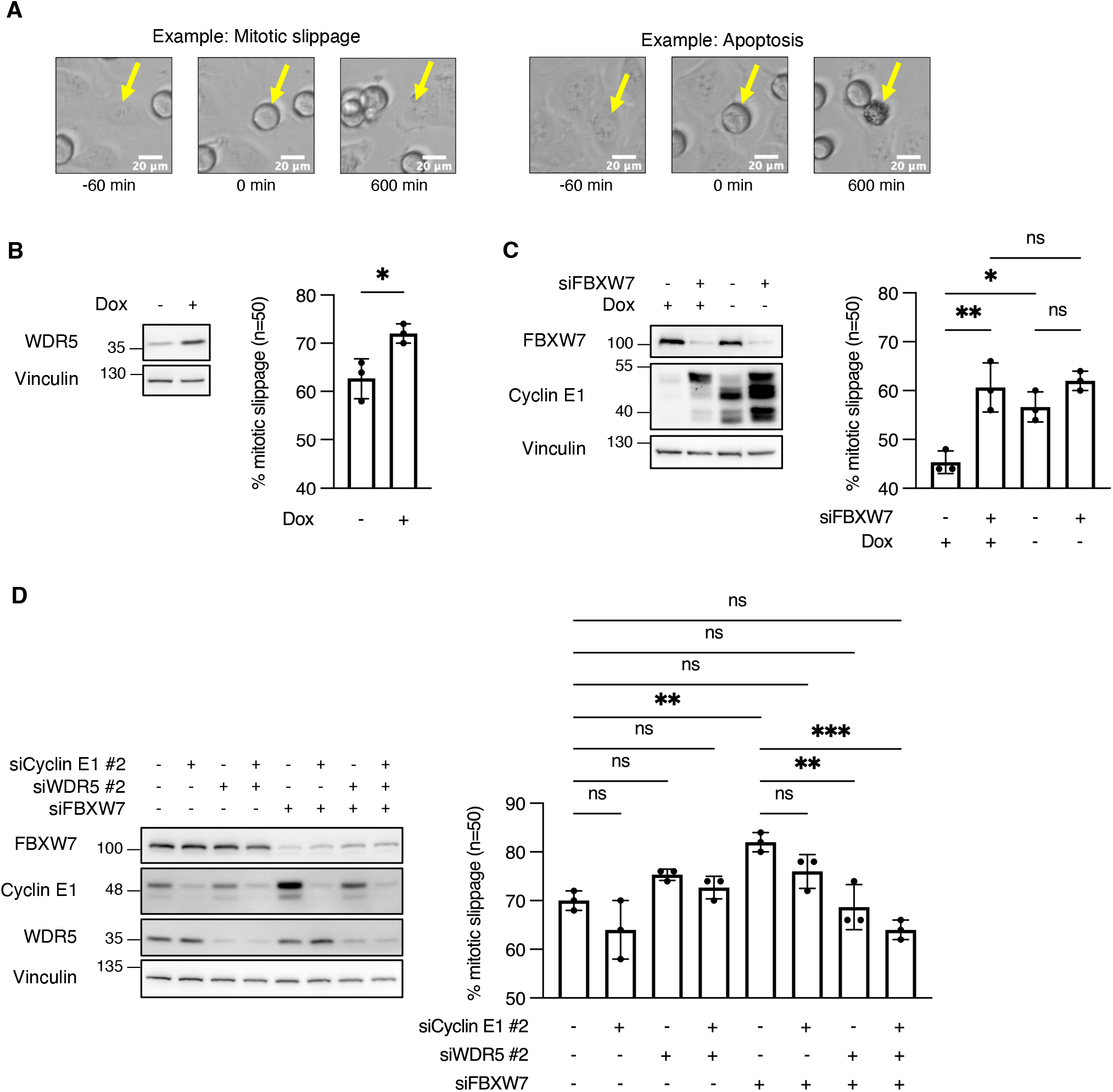
WDR5 and Cyclin E1 promote mitotic slippage. A, still images from live-cell imaging with U2OS T-Rex WDR5 showing different mitotic cell fates in response to 830 nM nocodazole. Arrows indicate cells undergoing mitotic arrest and slippage or apoptosis. B, U2OS T-Rex WDR5 were cultured in medium containing 2 µg/ml doxycycline for four days before the addition of 830 nM nocodazole and live-cell imaging. Mitotic cell fate of n = 50 cells was determined for each experiment. * p < 0.05, two-way unpaired students t-test, n = 3. C, U2OS tet-Off Cyclin E1 were cultured with or without 2 µg/ml doxycycline for four days. At day two and three, cells were transfected with 30 nM siRNA targeting Gl2 of FBXW7, as indicated. 48 h after the second transfection, 830 nM nocodazole were added and mitotic cell fates of n = 50 cells determined by live-cell imaging. ** p < 0.01, * p < 0.05, ns p > 0.05, one-way ANOVA with Tukey post-hoc test, n = 3. D, U2OS were transfected twice with 30 nM siRNA targeting Gl2, FBXW7, Cyclin E1 or WDR5, as indicated. 830 nM nocodazole were added 48 h after the second transfection and mitotic cell fates of n = 50 cells determined by live-cell imaging. *** p < 0.001, ** p < 0.01, p < 0.05, ns p > 0.05, one-way ANOVA with Tukey post-hoc test, n = 3.

Another key substrate of FBXW7 is Cyclin E1. Previous work has shown that overexpression of Cyclin E1 leads to accumulation of cells in early mitosis and chromosomal instability (29) (30). In addition, overexpression of Cyclin E1 has been suggested to weaken nocodazole-dependent mitotic arrest (31). Therefore, we asked whether overexpression of Cyclin E1 leads increased mitotic slippage and less cell death in response to nocodazole. We used an U2OS cell line where Cyclin E1 can be ectopically expressed after removal of doxycycline and performed live cell imaging experiments (32). We observed that overexpression of Cyclin E1 led to an increase in mitotic slippage to a similar extent as FBXW7 depletion. Simultaneous overexpression of Cyclin E1 and FBXW7 depletion did not further increase slippage (Fig. 5C). This data suggests that mitotic slippage could be promoted by ectopic expression of single FBXW7 substrates.

Next, we asked whether down-regulation of Cyclin E1 and WDR5 could rescue the mitotic slippage phenotype induced upon depletion of FBXW7. As shown in Fig. 5D, we find that the effect of FBXW7 depletion was partially rescued by knockdown of Cyclin E1 but more pronounced by the depletion of WDR5 or the combined depletion of both proteins. Interestingly, knockdown of Cyclin E1 or WDR5 did not reduce mitotic slippage in comparison to the siRNA control. We also showed that KMT2D depletion did not affect mitotic cell fate (Fig. 1C), suggesting that WDR5 exerts this function in regulation of mitotic cell fate independent of KMT2D. Taken together, our results support that WDR5 and Cyclin E1 are involved in the FBXW7-dependent regulation of mitotic cell fate.

### WDR5 and Cyclin E1 are equally required for the formation of polyploidy in response to anti-microtubule drugs

Mitotic slippage leads to the formation of tetraploid cells and further to polyploidy, which is reflected by an increase in DNA content. There is evidence for an involvement of Cyclin E1 and Aurora A in the formation of polyploidy after treating FBXW7 depleted cells with anti-microtubule drugs (4). Therefore, we asked whether WDR5 also plays a role in drug-induced polyploidy. We performed a FACS analysis of the DNA content in cells arrested in mitosis by addition of the microtubule stabilizing drug Taxol. HCT116 FBXW7 KO cells were treated with control, Cyclin E1, WDR5 or KMT2D specific siRNAs, respectively, and further treated following the scheme shown in Fig. 6A. Verification of increased WDR5 and CyclinE1 levels in response to FBXW7 knockout and reduction of target proteins by siRNA treatment is demonstrated in Fig. 6B. As previously shown (4), downregulation of FBXW7 leads to an increase of DNA content >4N in the presence of anti-microtubule drugs. Upon downregulation of WDR5 or Cyclin E1 in FBXW7 KO HCT116, respectively, the increase in DNA content could be markedly reduced (Fig. 6C), suggesting that WDR5 and Cyclin E1 function are required for formation of polyploid cells. Co-depletion of WDR5 and CyclinE1 did not further decrease the DNA-content (Fig. 6D). This could hint that both proteins support polyploidization through similar mechanisms. Depletion of KMT2D in HCT116 FBXW7 KO cells did not lead to a decrease in DNA content. Similar results were obtained when vincristine, a microtubule-destabilizing drug, was used instead of Taxol to arrest cells in mitosis (Fig. S2). Taken together, these data provide evidence that WDR5 and Cyclin E1 are equally required for the polyploidization of FBXW7-depleted cells in response to different anti-microtubule drugs.

**Figure 6.**
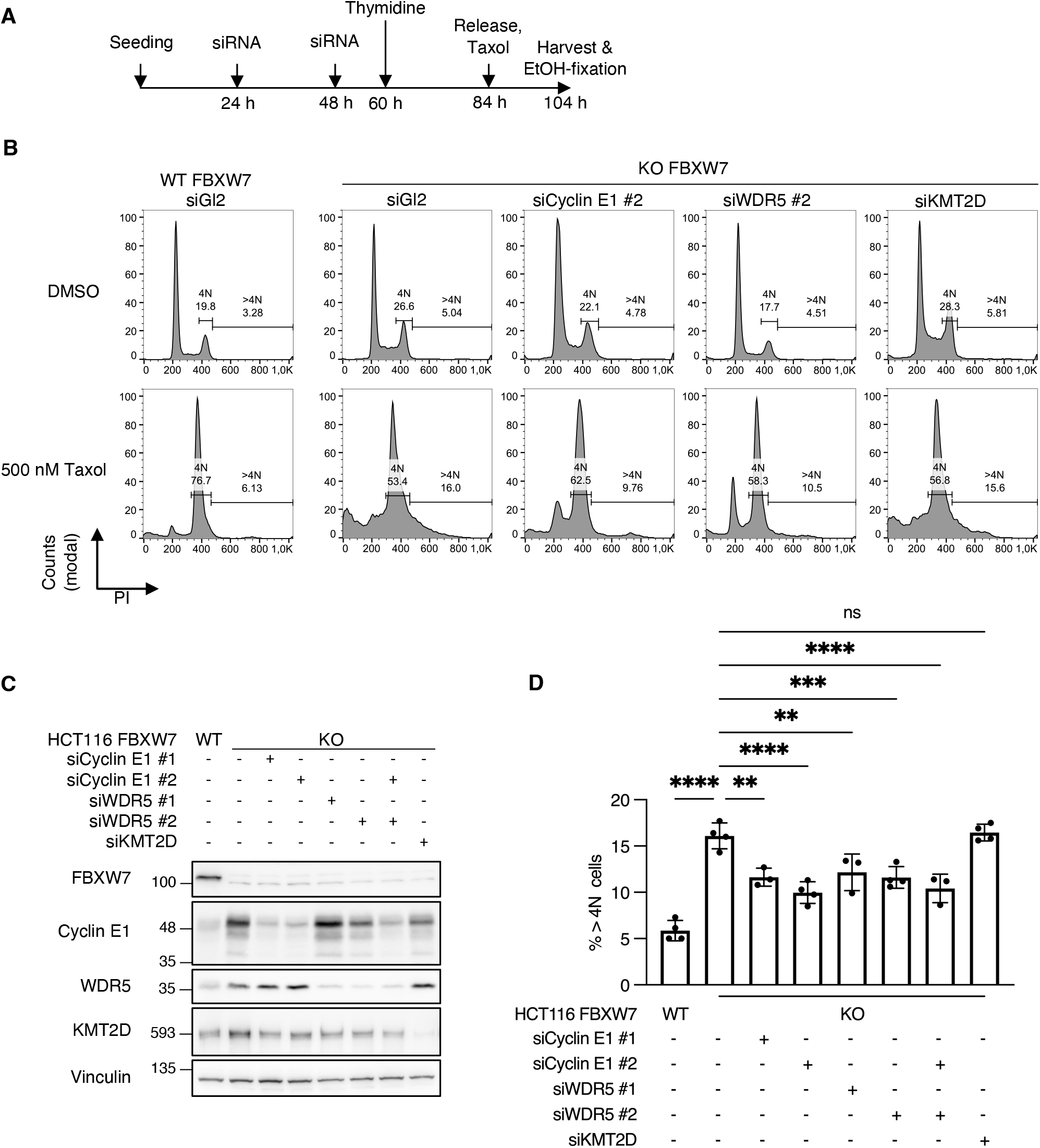
WDR5 and Cyclin E1 are equally required for drug-induced polyploidy in response to anti-microtubule drugs. A, schematic depicting the protocol followed to determine drug-induced polyploidy of HCT116. HCT116 WT and FBXW7 knockout were transfected twice with 30 nM siRNA and blocked in S-Phase with 2 mM Thymidine for 24 h. Cells were released into medium containing 500 nM Taxol and harvested and fixed after 20 h. B, exemplary histograms showing DNA contents of HCT116 determined by FACS-Scan analysis. HCT116 were treated following the protocol in A, and DNA stained with PI for 15 min at RT. Histograms are show modal counts and fraction of G2/M (4N) and polyploid (>4N) cells. C, HCT116 were transfected following the protocol from A but without synchronization and Taxol treatment. Asynchronous cells were harvested 48 h after the second siRNA transfection. D, Statistical analysis of B. **** p < 0.0001, *** p < 0.001, ** p < 0.01, ns p > 0.05, one-way ANOVA with Tukey post-hoc test, n = 3-4.

## Discussion

WDR5 localizes to the spindle microtubules during mitosis and its depletion leads to chromosome misalignment and spindle assembly defects (19). Here we report a new function of WDR5 in mitotic cell fate. We not only found that WDR5 plays a role in mitotic slippage caused by prolonged mitotic arrest in response to treatment with anti-microtubule drugs but also that WDR5 is a substrate and binding partner of the tumor suppressor FBXW7 which is involved in mitotic cell fate decision (4) (5) (11). High levels of WDR5 induced by absence of FBXW7 promote mitotic slippage. In the future, it would be interesting to find out whether FBXW7-mediated degradation of WDR5 is also involved in canonical functions of WDR5, for instance histone methylation (H3K4).

Anti-microtubule drugs arrest or delay cells in mitosis due to the activation of the mitotic or spindle assembly checkpoint (6). If the SAC cannot be satisfied anymore, mitosis remains uncompleted as cells either undergo cell death or perform mitotic slippage (11). MCL-1, a member of the anti-apoptotic Bcl-2 family of proteins that suppress the activation of caspases is degraded by ubiquitin mediated degradation (5,9,33).

The tumor suppressor FBXW7 has been also implicated in cell fate decision but it is not yet clear whether it plays a role in regulating MCL-1 protein levels (5,10,11). Therefore, it is important to identify and characterize substrates of FBXW7 involved in mitotic cell fate. Similar to the regulation of KMT2D by FBXW7 (22), we show that WDR5 protein levels are regulated by FBXW7 through ubiquitin-mediated degradation suggesting that two proteins of the MLL4 complex are substrates of FBXW7. However, while WDR5 upregulation promotes mitotic slippage, KMT2D upregulation does not affect exit from drug-induced mitotic arrest suggesting that the deregulation of both proteins affects cancer cells differently. In addition, binding of FBXW7 to WDR5 does not require KMT2D (Fig. 2D). Together our data implicate that FBXW7 targets WDR5 for degradation and that WDR5 upregulation supports chemoresistance of FBXW7-depleted cancer cells by mitotic slippage while the FBXW7/KMT2D axis was previously shown to function in oxidative phosphorylation in B-cell malignancies (22). As the overexpression of each of the two FBXW7 substrates, WDR5 and Cyclin E1, induced mitotic slippage, we suggest that both substrates work downstream of FBXW7 in mitotic slippage induction but whether they act in the same pathway remains unclear. In addition, we cannot exclude that other FBXW7 substrates could also function in mitotic slippage. For example, c-Myc, another key substrate of FBXW7, promotes cell death following prolonged mitotic arrest (34).

In summary, our results open the venue for development of drugs to prevent mitotic slippage in response to treatment with anti-microtubule drugs. WDR5 has been characterized as a target for the development of small molecule inhibitors (35) (36). Future work could involve testing of WDR5 inhibitors in FBXW7-deficient or mutated cancer types.

## Experimental procedures

### Cell lines and cell culture

HEK293T (ACC 635; DSMZ, Braunschweig, Germany), HeLa (ATCC CCL-2), HCT116 and HCT116 KO FBXW7 (B. Vogelstein, Johns Hopkins University, Baltimore MD, USA), U2OS (ATCC HTB-96), U2OS Flp-In-T-rex (J.D. Pravin, Ohio State University) and U2OS Tet-Off Cyclin E1 (J. Bartek, Karolinska Institute, Stockholm, Sweden) were maintained in Dulbecco’s Modified Eagle’s Medium containing 4.5 g/l glucose (Gibco 41965-039). For U2OS Tet-Off Cyclin E1, 2 µg/ml Doxycycline (Sigma D-9891) were added to the cell culture medium. DLD1 and DLD1 KO FBXW7 (B. Vogelstein, Johns Hopkins University, Baltimore MD, USA) were cultivated in RPMI1640 (Sigma R8758). All media were supplemented with 10% FBS (Gibco 10270-106) and 1% penicillin/streptomycin (Sigma P0781). Cell lines were grown in a humidified CO_2_ incubator at 37°C and 5% CO_2_. U2OS Flp-In-T-Rex WDR5 and WDR5 F133A stable cell lines were generated following the manufacturer’s instructions (Life Technologies). Protein expression was induced by addition of 2 µg/ml Doxycycline for at least 72 h. GSK3β activity was blocked with 5 µM CHIR-99021 (MedChemExpress, HY-10182) and cullin-RING E3 ubiquitin ligases with 5 µM MLN4924 (Cell Signaling Technology, 85923) for 5 h prior to cell harvest.

### Plasmids, cloning, and mutagenesis

Human FBXW7α and WDR5, cDNAs were obtained from the Genomics and Proteomics Core Facility at the DKFZ (Heidelberg, Germany). Full-length FBXW7α was cloned into pCMV-3Tag1C and pEGFP-C1 via EcoRI and HindIII sites. WDR5 cDNA was subcloned to pCMV-3TagA using HindIII and XhoI, to pEGFP-C2 with ApaI and EcoRI and to pGEX-4T1 with EcoRI and XhoI. pCDNA3-Flag-FBXW7α and pCDNA3-Flag-FBXW7α T439I, S642A, T463A, R465A mutant were a gift from Michele Pagano (NYU Grossmann School of Medicine, New York). pCVM-Tag5A-Myc-GSK3β was a gift from Mien-Chie-Hung (37) (Addgene #16260). Mutants of cDNA were generated by site-directed mutagenesis. For the generation of stable WDR5 expressing cell lines using the T-Rex system (WDR5 constructs were subcloned to pcDNA5-FRT-TO using HindIII and XhoI.

### Transient plasmid and siRNA transfections

HEK-293T cells were transiently transfected with plasmid DNA using polyethylenimine (Polysciences) at a ratio of DNA 1:3.4 polyethylenimine. Proteins were overexpressed for 24-48 h. Lipofectamine 2000 (Invitrogen) was used to transfect DLD1 cell lines with plasmid DNA following the manufacturer’s instructions. Transfection mixes were removed 6 h after transfection and fresh growth medium was added.

Lipofectamine 2000 was used for transfection of HeLa or U2OS, and Lipofectamine RNAiMAX (Invitrogen) was used for transfection of HCT116 with siRNA. Cells were transfected with 30 nM of siRNA 24 h and 48 h after seeding and further cultivated for at least 48 h after the second siRNA transfection.

The following siRNA sequences were used:

Gl2 (firefly luciferase), 5’-CGUACGCGGAAUACUUCGAdTdT-3’;

pan-FBXW7, 5’-ACAGGACAGUGUUUACAAAdTdT-3’ (38)

WDR5 #1, 5’-UUAGCAGUCACUCUUCCACUUdTdT-3’ (19)

WDR5 #2, 5’-GCUCAGAGGAUAACCUUGUdTdT-3’ (39)

Cyclin E1 #1, 5’-CCUCCAAAGUUGCACCAGUUUdTdT-3’ (40);

Cyclin E1 #2, 5’-UGACUUACAUGAAGUGCUAdTdT-3’

siGSK3β #1, 5’-GAAGUCAGCUAUACAGACAdTdT-3’

siGSK3β #2, 5’-GGUCACGUUUGGAAAGAAUdTdT-3’ (41)

siKMT2D, 5’-CCCACCUGAAUCAUCACCUdTdT-3’ (42)

### Cell lysis, coimmunoprecipitation and western blot analysis

For Western blot analysis, cells were harvested and washed with ice-cold PBS. Cell pellets were lysed in 3-5 volumes of NP40 or RIPA buffer. NP40 buffer (40 mM Tris-HCl pH 7.5, 150 mM NaCl, 5 mM EDTA, 10 mM β-glycerophosphate, 5 mM NaF, 0.5% NP40, 1 mM DTT, 10 µg/ml TPCK, 5 µg/ml TLCK, 0.1 mM NA_3_VO_4_, 1 µg/ml Aprotinin, 1 µg/ml Leupeptin, 10 µg/ml Trypsin inhibitor from soybean) was used for all immunoprecipitations. For all other samples, RIPA buffer (50 mM Tris-HCl pH 7.4, 1% NP40, 0.5% Na-deoxycholate, 0.1% SDS, 150 mM NaCl, 2 mM EDTA, 50 mM NaF, 1 mM DTT, 10 µg/ml TPCK, 5 µg/ml TLCK, 0.1 mM NA_3_VO_4_, 1 µg/ml Aprotinin, 1 µg/ml Leupeptin, 10 µg/ml Trypsin inhibitor from soybean) was used. Cell lysates were incubated on ice for 30 min and cleared by centrifugation at 16.100x g for 20 min. For SDS-PAGE, cell extracts were mixed with 2x Laemmli buffer or 2x LDS sample buffer (samples for KMT2D analysis) and incubated at 95 °C for 5 min or 72 °C for 10 min, respectively. Immunoprecipitations were performed as described previously (8). Proteins were resolved by SDS-PAGE and transferred to nitrocellulose membranes. For the analysis of KMT2D protein levels, a TRIS-acetate gel and buffer system was used (43). Proteins were detected using following antibodies: α-FBXW7α (A301-720), α-Rbbp5 (A300-109A), α-KMT2D (A300-BL1185), α-MLL1 (A300-374A) and α-SETD1A (A300-289A) antibodies were obtained from Bethyl. α-WDR5 (G9), α-Cyclin E1 (HE12) and α-Myc (9E10) antibodies were purchased from Santa Cruz Biotechnology Inc., α-Flag-tag (F3165), α-Vinculin (V9131) and α-Tubulin (T6074) antibodies were from Sigma. α-c-Myc (9402) antibody was from Cell Signaling Technology, α-HA-tag (16B12) antibody from Babco and the α-GFP antibody was produced in-house.

### GST pull-down assay

Recombinant GST-WDR5 was expressed in *E. coli* Rosetta (DE3) and purified using glutathione CL-4B Sepharose (Sigma 49739). His-FBXW7/Skp1 was a gift from Frauke Melchior (University of Heidelberg, Germany). For *in-vitro* GST pull-down assays, 10 µg of purified GST-WDR5 and 10 µg of His-FBXW7/Skp1 were mixed in 200 µl NP40 buffer and incubated on a rotating wheel at 4 °C for 1 h. 10 µL glutathione Sepharose beads were resuspended in 200 µL NP40 buffer, added to the mixture and incubated at 4 °C for 2 h. Beads were washed four times with NP40 buffer and proteins eluted by incubation with 25 µL 2x Laemmli buffer at 95 °C for 5 min.

Cycloheximide chase

For the analysis of WDR5 protein stability, DLD1 WT and DLD1 KO FBXW7 were transfected with pCMV-3Tag1A-WDR5 using Lipofectamine 2000. 24 h after transfection, 300 µg/ml cycloheximide (ChemCruz, Santa Cruz Biotechnology, SC-3508) were added to inhibit protein synthesis. Samples were harvested at different time points for up to 24 h. Protein degradation via the 26S proteasome was blocked by adding 10 µM MG132 (Sigma-Aldrich, C2211) 3 h before the addition of cycloheximide. 10 µM MG132 were then also added to the medium with cycloheximide.

### Ubiquitination assays

HEK293T were transiently co-transfected with the indicated plasmids. 24 h after transfection, cells were treated with 10 µM MG132 for 5 h. GSK3β or cullin-RING E3 ligase activity was blocked by addition of 5 µM CHIR-99021 or 5 µM MLN4924, respectively, for 4 h prior to addition of MG132. Cells were collected and lysed in complete NP40 buffer with 20 mM N-ethylmaleimide (Sigma, E3876) and Flag-immunoprecipitations performed as described above. Samples of whole cell extracts and eluates were denatured with LDS sample buffer for 10 min at 72°C. Proteins were resolved by SDS-PAGE and detected after Western blotting.

### Mass spectrometry based SCF-FBXW7 substrate screen

Eluates from α-Flag-FBXW7 and Flag-FBXW7 WD40 mutant (T439I, S642A, T463A, R465A) immunoprecipitations were analyzed by SDS-PAGE and stained by Colloidal Coomassie. Mass spectrometry and data processing was performed by the EMBL Proteomics Core Facility Heidelberg. Briefly, whole gel lanes were cut and proteins reduced with 10 mM DTT in 100 mM NH_4_HCO_3_ for 30 min at 56 °C. 180 µl acetonitrile were added at room temperature for 15 min, followed by replacement with 200 µL of 55 mM chloroacetamide in 100 mM NH_4_HCO_3_ and alkylation for 20 min in the dark. Gel pieces were washed twice with acetonitrile at room temperature for 15. Proteins were in-gel digested by addition of 200 µL of 2 ng/µl trypsin (Promega, V511A) in resuspension buffer (Promega, V542A) and incubation on ice for 30 min and then at 37 °C overnight. Peptides were extracted with 50% acetonitrile and 1% formic acid, dried and reconstituted in 0.1% (v/v) formic acid. After a reverse phase cleanup step, peptides were reconstituted in 10 µL of 100 mM Hepes/NaOH, pH 8.5 and reacted with 80 µg of TMT10plex (Thermo Scientific, 90111) dissolved in 4 µL acetonitrile for 1 h at room temperature. Peptides were mixed 1:1, subjected to reverse phase clean-up step and analyzed by LC-MS/MS on a Q Exactive Plus (Thermo Scientific) as previously described (44). Acquired data were analyzed using IsobarQuant and Mascot V2.4 (Matrix Science). Peptide mass error tolerance was 10 ppm for full scan MS spectra and 0.02 Da for MS/MS spectra and a false discovery rate below 0.01 was required on peptide and protein level. Two replicates were performed for each condition and results compared using the LIMMA package for R.

### Quantitative real-time PCR

Total RNA was extracted using TRIzol ™ reagent (Invitrogen) following the manufacturer’s instructions. Potential genomic DNA contaminants were digested with DNAse (Sigma, AMPD1) and first strand cDNA was synthesized with RevertAid cDNA Synthesis Kit (ThermoScientific, K1621). Quantitative real-time PCR was performed using Power SYBR^®^ Green PCR Master Mix (Applied Biosystems, 4368577) and PCRs were performed in a QuantStudio 5 cycler (Applied Biosystems). Relative mRNA expression was calculated using the ΔΔCT method.

WDR5 forward 5’-ATGCGACAGAGACCATCATAG-3’

WDR5 reverse 5’-CGTGAGGATATGGGATGTGAA-3’

GAPDH forward 5’-CAAGGCTGAGAACGGGAAG-3’

GAPDH reverse 5’-TGAAGACGCCAGTGGACTC-3’

### Live-cell imaging

Mitotic cell fates were analyzed by live-cell imaging. U2OS were transfected with 30 nM siRNA 24 h and 48 h after seeding. 24 h after second transfection, cells were seeded to 8-well imaging dishes (Sarstedt, 94.6170.802) at 50% confluency. For U2OS T-Rex WDR5 clones and U2OS tet-Off Cyclin E1, subcultures were cultivated in medium with or without 2 µg/ml Doxycycline for 72 h before seeding to imaging dishes. In addition, U2OS tet-Off Cyclin E1 were transfected twice with 30 nM siRNA. 24 h after seeding, 830 nM nocodazole were added to all wells and the dish was placed in a microscopy incubation chamber at 5% CO_2_ and 37 °C. Cells undergoing mitotic arrest were monitored using a Zeiss Cell Observer Z1 inverted microscope and a 10x /0.3 EC PlnN Ph1 DICI objective. Phase-contrast images of multiple positions were taken every 10 min for up to 60 h using the Zeiss ZEN blue software. Cell fates during mitotic arrest were analyzed with ImageJ Fiji.

### Flow cytometry analysis

DNA content of HCT116 and HCT116 KO FBXW7 was determined by DNA staining of EtOH-fixed cells with 30 µg/ml propidium iodide and flow cytometric analysis. DNA content was determined on a FACSCalibur (BD) and raw data analyzed using FlowJo v10 (BD).

### Statistical analysis

All statistical analysis, if not stated otherwise, were performed with GraphPad Prism v9. Quantitative data was collected from at least three independent experiments and are represented as mean ± SD. Statistical significance was analyzed by unpaired, two-tailed, Student’s t test with Welch’s correction or one-way ANOVA with Tukey post-hoc test, as indicated in each figure legend. p values of less than 0.05 were considered statistically significant (ns p > 0.05, * p < 0.05, ** p < 0.01, *** p < 0.001, **** p < 0.0001).

## Figure legends

**Figure S1**.

A, Coomassie staining showing samples of Flag-FBXW7 immunoprecipitations designated for mass spectrometric analysis. Flag-FBXW7 constructs were overexpressed in HEK293T cells and α-Flag immunoprecipitations performed using α-Flag M2 beads. Proteins were separated by SDS-PAGE and staining using colloidal Coomassie. M – Marker, 1 – Flag-EV, 2 – Flag-FBXW7, 3 – Flag-FBXW7 T439I, S462A, T463A, R465A.

B, Summary of hits from Flag-FBXW7 IP-MS. Significance between Flag-FBXW7 and Flag-FBXW7 WD40 mutant interactors was determined via t-statistics using LIMMA R-package; proteins were considered significant if they showed a fold change of at least 50% and an adjusted p-value after Benjamini and Hochberg below 5%, n = 2.

**Figure S2**.

Statistical analysis of B showing a significant reduction of drug-induced polyploidy of HCT116 FBXW7 KO by depletion of Cyclin E1 or WDR5. **** p < 0.0001, *** p < 0.001, ** p < 0.01, * p < 0.05, ns p > 0.05, one-way ANOVA with Tukey post-hoc test, n = 3-4

The MS proteomics data have been deposited to the ProteomeXchange Consortium (http://proteomecentral.proteomexchange.org) via the PRIDE partner repository with the data set identifier 1-20220720-46266.

Login details:

Password: FBXW7F045WDR5

## Acknowledgements

We thank J. Bartek, F. Melchior, M. Pagano and B. Vogelstein for providing reagents and cell lines. We acknowledge the Microscopy Core Facility for providing equipment and technical assistance. We thank A. Turi da Fonte Dias for comments on the manuscript.

This work was supported by a grant from the Wilhelm Sander-Stiftung Grant No. 2019.049.1.

## Notes

### Competing Interest Statement

The authors have declared no competing interest.

