## Supplemental Figure 1 for "FBXW7 triggers degradation of WDR5 to prevent mitotic slippage"

**A**

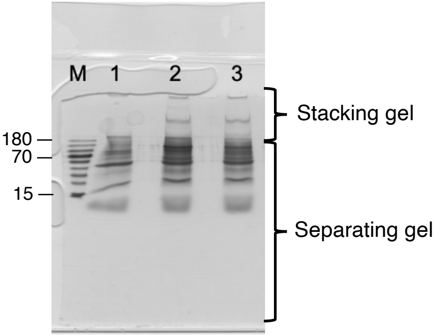


**B**

| gene | top3 | qupm | logFC | AveExpr | t | pvalue.limma | fdr.limma | B |
|---|---|---|---|---|---|---|---|---|
| TP53 | 8,24802 | 21 | 2,14202867 | 28,6416835 | 18,8849177 | 1,5937E-08 | 9,865E-06 | 10,1466484 |
| KMT2D | 6,96643 | 18 | 1,03086358 | 27,6379627 | 10,1693565 | 3,2198E-06 | 0,00075313 | 5,13607199 |
| PAXIP1 | 6,89502 | 2 | 1,02405246 | 25,9125603 | 10,0098786 | 3,6712E-06 | 0,00075313 | 5,00338737 |
| MYC | 6,39471 | 3 | 1,0734975 | 24,8920443 | 9,6743658 | 4,8668E-06 | 0,00075313 | 4,71732556 |
| KDM6A | 6,49671 | 3 | 1,18532286 | 24,1465563 | 8,41896575 | 1,5107E-05 | 0,00187027 | 3,5565237 |
| PRR36 | 6,48068 | 5 | 1,34787304 | 25,4154348 | 7,60063293 | 3,4082E-05 | 0,00351612 | 2,71364703 |
| WDR5 | 6,93304 | 5 | 0,73760404 | 26,253807 | 7,18073732 | 5,3118E-05 | 0,00469719 | 2,25149011 |
| ASH2L | 6,59014 | 3 | 0,67972041 | 25,4975787 | 6,86494596 | 7,5114E-05 | 0,00581194 | 1,88969623 |

Supplemental figure 1
