## Supplementary figures and images for "FBXW7 triggers degradation of WDR5 to prevent mitotic slippage"

### Supplemental Figure 2

A

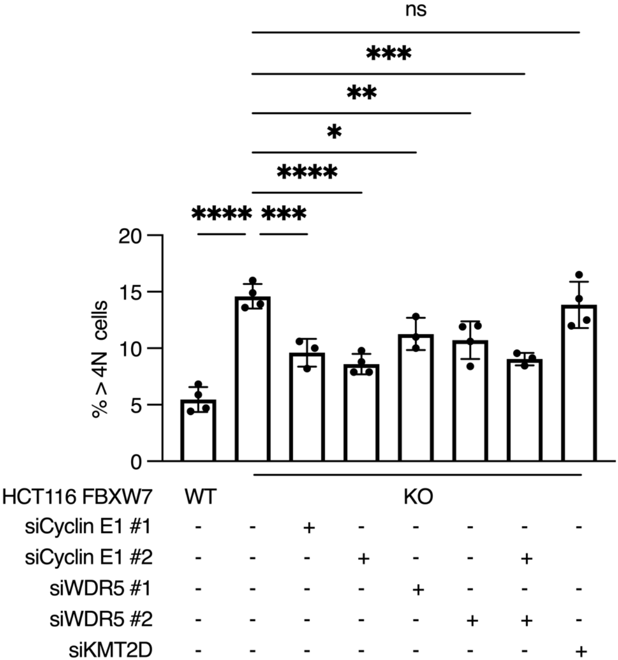


Supplemental figure 2
